# Population-specific brain charts reveal Chinese-Western differences in neurodevelopmental trajectories

**DOI:** 10.1101/2025.06.17.659820

**Authors:** Lianglong Sun, Wen Qin, Xinyuan Liang, Caihong Wang, Weiwei Men, Yunyun Duan, Xue-Ru Fan, Qing Cai, Shijun Qiu, Meiyun Wang, Qiyong Gong, Yanghua Tian, Peipeng Liang, Zeyu Liu, Xiaochu Zhang, Hongwen Song, Zhaoxiang Ye, Peng Zhang, Qi Dong, Sha Tao, Wenzhen Zhu, Jintao Zhang, Fang Xie, Jianfeng Feng, Jing Zhang, Chao Liu, Qiujin Qian, Bing Zhang, Ming Meng, Li Hu, Jia-Hong Gao, Tianzi Jiang, Xiongzhao Zhu, Yuhan Zhang, Liping Liu, Hanjun Liu, Weihua Liao, Dawei Wang, Huali Wang, Tengfei Guo, Zhengjia Dai, Su Lui, Kai Xu, Lingjiang Li, Peng Xie, Chunliang Feng, Guangbin Cui, Jinsong Wu, Xuntao Yin, Guosheng Ding, Junfang Xian, Lianping Zhao, Jie Lu, Zhifen Liu, Ying Han, Zhen Yuan, Xilin Zhang, Tianmei Si, Fuqing Zhou, Yanchao Bi, Dan Wu, Fei Gao, Fei Wang, Shaozheng Qin, Gang Wang, Feng Chen, Zhiqiang Zhang, Jing Sui, Huafu Chen, Jinhua Cai, Shuwei Liu, Zuojun Geng, Chen Zhang, Ning Mao, Hong Yin, Bo Liu, Heng Ma, Bo Gao, Yanwei Miao, Xiang-Zhen Kong, Yuan Zhou, Li Liu, Jianping Hu, Liang Wang, Quan Zhang, Hua Shu, Peijun Wang, Tatia M. C. Lee, Qingjiu Cao, Li Yang, Xi Zhang, Wenbo Luo, Meng Liang, Hongxiang Yao, Meng Li, Hao Huang, Yun Peng, Zaizhu Han, Chao Zhou, Haibo Xu, Ming Feng, Wen Shen, Yuzheng Hu, Huajun Chen, Ying Wang, Gaolang Gong, Zhihan Yan, Xiaojun Xu, Jun Liu, Guangxiang Chen, Pan Wang, Yunjun Yang, Dezhong Yao, Tong Han, Huiguang He, Ce Chen, Qihong Zou, Hesheng Liu, Hui Zhang, Chao Chai, Chunming Lu, Yiheng Tu, Yong Liu, Danhua Lin, Weihua Zhao, Xiufeng Xu, Xiaoli Liu, Zaixu Cui, Zheng Wang, Ruiwang Huang, Zhanjiang Li, Yunzhe Liu, Xiaojun Li, Xiujie Yang, Nan Zhang, Antao Chen, Bin Zhang, Pengmin Qin, Chen Liu, Zhenwei Yao, Yanjun Wei, Huishu Yuan, Feng Wang, Yu Zhang, Quan Zhang, Fang Hu, Huan Xie, Xuehai Wu, Jiaojian Wang, Guoguang Fan, Zhiqun Wang, Dongling Zhang, Hui Zhong, Yonggang Wang, Lijun Bai, Yongmei Li, Xinhua Wei, Jinhui Wang, Yi Zhang, Hongjian He, Shuyu Li, Tijiang Zhang, Fan Jiang, Jian Yang, Feiyan Chen, Feng Liu, Huaigui Liu, Nan Chen, Jinzhu Yang, Bo Hou, Chu-Chung Huang, Jiajia Zhu, Huanhuan Cai, Dongtao Wei, Qunlin Chen, Ying Wei, Peifang Miao, Yunxia Li, Yaou Liu, Ning Yang, Xiaoxue Gao, Yujie Liu, Yu Shen, Xiaoqi Huang, Gong-Jun Ji, Alzheimer’s Disease Neuroimaging Initiative, CHIMGEN Consortium, DIDA-MDD Working Group, MCADI, Chinese Lifespan Brain Mapping Consortium, Longjiang Zhang, Jiang Qiu, Yongqiang Yu, Ching-Po Lin, Feng Feng, Kuncheng Li, Chunshui Yu, Yong He

## Abstract

Human brain charts provide unprecedented opportunities for decoding neurodevelopmental milestones and establishing clinical benchmarks for precision brain medicine ^1-7^. However, current lifespan brain charts are primarily derived from European and North American cohorts, with Asian populations severely underrepresented. Here, we present the first population-specific brain charts for China, developed through the Chinese Lifespan Brain Mapping Consortium (Phase I) using neuroimaging data from 43,037 participants (aged 0−100 years) across 384 sites nationwide. We establish the lifespan normative trajectories for 296 structural brain phenotypes, encompassing global, subcortical, and cortical measures. Cross-population comparisons with Western brain charts (based on data from 56,339 participants aged 0−100 years) reveal distinct neurodevelopmental patterns in the Chinese population, including prolonged cortical and subcortical maturation, accelerated cerebellar growth, and earlier development of sensorimotor regions relative to paralimbic regions. Crucially, these Chinese-specific charts outperform Western-derived models in predicting healthy brain phenotypes and detecting pathological deviations in Chinese clinical cohorts. These findings highlight the urgent need for diverse, population-representative brain charts to advance equitable precision neuroscience and improve clinical validity across populations.

## Introduction

Human brain charts across the entire lifespan are indispensable for advancing both basic neuroscience and clinical research in neuropsychiatric disorders. The continuous aggregation of large-scale neuroimaging datasets has enabled the construction of population-level structural ^1-3^ and functional ^4, 5^ brain charts. These transformative resources are reshaping our understanding of neurodevelopmental milestones and providing critical clinical benchmarks for identifying pathological deviations in brain disorders ^6, 7^. However, the currently available lifespan brain charts are predominantly derived from European and North American cohorts (i.e., Western-centric models), with a severe underrepresentation of Asian populations ^1-4^. The largest lifespan brain chart study to date ^1^ (involving 101,457 individuals across 286 global sites) included fewer than 4,500 participants (< 4%) from East Asian ancestry, with a strikingly low representation from China (fewer than 3000 participants, < 3%, from 8 sites in only 2 cities), which is a nation constituting 18% of the world’s population. This representative bias raises critical questions about the universality of current brain charts and their clinical utility across ancestral populations.

Compelling evidence demonstrates that, compared with individuals of European ancestry, Chinese individuals exhibit distinct neuroanatomical features ^8-10^ and developmental trajectories ^11-14^, particularly in the frontoparietal cortices. These inter-ancestry differences, shaped by complex gene-culture-environment interactions ^15-19^, fundamentally challenge the generalizability of brain charts derived predominantly from Western populations to the Chinese population. Supporting this, recent studies have revealed ethnic disparities (white/Asian/black) in deviation scores within the UK Biobank cohort ^20^ and reduced accuracy in cross-population brain-behavior prediction ^21^. Together, these empirical failures challenge the presumption of neurodevelopmental universality across ethnic groups, necessitating the urgent establishment of population-specific reference frameworks through large-scale, ethnically representative neuroimaging datasets to ensure both scientific rigor and accurate clinical assessment.

To address this gap, the Chinese Lifespan Brain Mapping (C-LBM) Consortium presented its Phase I efforts to establish the first population-specific brain growth charts for Chinese individuals, based on a large-scale neuroimaging dataset of 43,037 participants (0−100 years of age, 384 sites across 29 provinces; Fig. 1a, b). By implementing rigorous quality control, harmonized analytical pipelines, and standardized statistical modelling, we delineate neurodevelopmental trajectories of 296 brain structural phenotypes in the Chinese population and conduct cross-population comparisons with Western cohorts (56,339 participants aged 0−100 years) (Fig. 1c-e). Our established Chinese brain charts (1) uncover neurodevelopmental milestones in previously under-explored brain phenotypes, (2) reveal robust and replicable cross-population differences in maturation timelines, (3) improve the phenotypic predictive accuracy for healthy Chinese samples, and (4) mitigate misestimation of brain deviations in Chinese neuropsychiatric conditions. These brain charts not only fill a critical knowledge gap in understanding population-specific milestones of brain maturation but also establish a blueprint for developing ethnically specific reference standards worldwide, thereby advancing equitable precision medicine in brain health.

**Figure 1.**
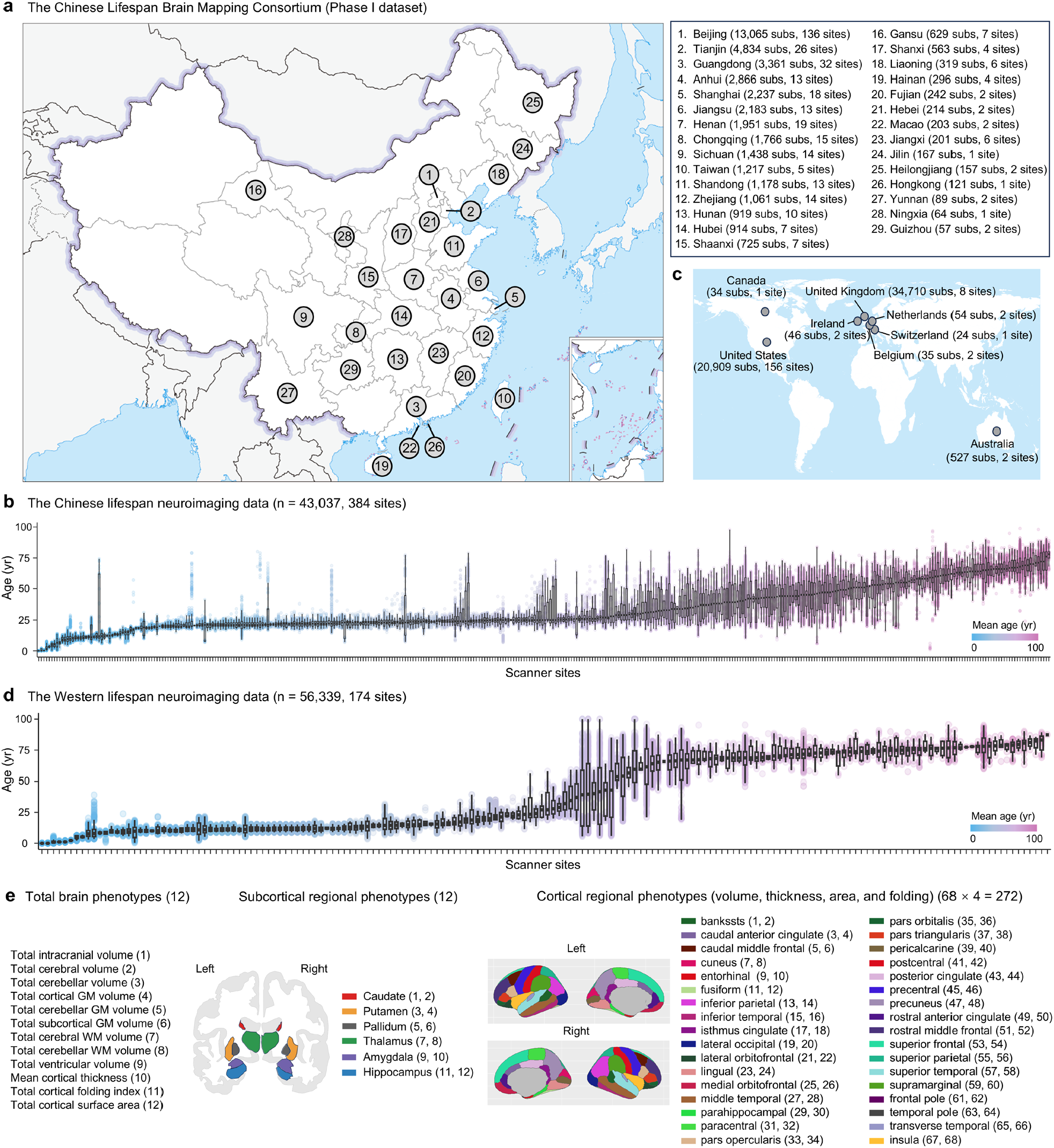
Structural neuroimaging data of Chinese and Western populations. **a**, Phase I dataset of the Chinese Lifespan Brain Mapping (C-LBM) Consortium, comprising neuroimaging data collected across 29 provinces in China. The right panel shows the number of participants and scanning sites within each province. **b**, Quality-controlled Chinese lifespan neuroimaging dataset derived from 384 scanning sites, comprising 43,037 healthy participants aged 0–100 years. **c**, Western neuroimaging data aggregated from eight countries: the United States, the United Kingdom, Canada, Ireland, the Netherlands, Belgium, Switzerland, and Australia. **d**, Quality-controlled Western lifespan neuroimaging dataset derived from 174 scanning sites, including 56,339 healthy participants aged 0–100 years. Box plots depict the age distribution of participants at each site. Boxes represent the interquartile range (25th–75th percentiles), with the median indicated by a horizontal line; whiskers extend to 1.5 times the interquartile range, and points beyond are plotted as outliers. **e**, Structural brain phenotypes used for normative model charts, including total brain phenotypes (n = 12), subcortical regional phenotypes (n = 12), and cortical regional phenotypes (n = 272). subs, subjects; yr, year.

## Results

Following rigorous quality control (detailed in the Methods section), we analysed high-quality brain structural MRI data from 99,376 healthy participants aged 0–100 years. The detailed demographics of the datasets are presented in Supplementary Table 1. Using C-LBM Phase I data (43,037 participants, 56.8% females, 384 sites in China), we first constructed sex-stratified Chinese lifespan normative models for 296 structural brain phenotypes. For cross-population comparisons, we developed Western-based models (56,339 participants, 50.3% female, 174 sites across Europe/North America/Oceania, excluding Asian ancestry). Following World Health Organization guidelines ^22^, we modelled nonlinear growth trajectories using generalized additive models for location, scale and shape (GAMLSS) ^22, 23^, incorporating the scanner as a random covariate. Comparative analyses revealed significant population-level neurodevelopmental divergences between the Chinese and Western models. Through extensive sensitivity analyses, including bootstrapping, age-stratified balanced resampling, leave-one-site-out analysis, split-half replication, imaging quality-controlled modelling, and ethnicity-controlled modeling, we confirmed the robustness and reproducibility of our population-specific lifespan models (for details, see Sensitivity Analyses). Using Chinese case-control cohorts (2,591 healthy controls; 2,865 patients), we further demonstrated the superior performance of our Chinese model in both predicting typical phenotypic neurodevelopment (out-of-sample accuracy) and identifying pathological phenotypic deviations compared with the Western-based models.

### Population-specific normative growth of global brain phenotypes

Our analysis of Chinese lifespan growth charts revealed distinct neurodevelopmental timelines across global brain phenotypes. The volumetric metrics (excluding ventricles) followed inverted U-shaped curves, with phenotype-specific peak maturation (Fig. 2). Specifically, the total intracranial volume peaked during early adulthood (19.9 years, 95% confidence interval (CI) 19.3–20.4), whereas the total cerebral (13.4 years, 95% CI 12.9–14.0) and cerebellar (15.3 years, 95% CI 14.6–16.0) volumes reached their maxima earlier in adolescence (Fig. 2a-c). Grey matter volume (GMV) exhibited sequential maturation from childhood to adolescence: cortical (6.5 years, 95% CI 6.2–6.7), cerebellar (11.9 years, 95% CI 11.3–12.5), and subcortical (16.6 years, 95% CI 16.1–17.1) (Fig. 2d-f), while white matter volume (WMV) development extended into adulthood (cerebral: 31.0 years, 95% CI 30.6–31.5; cerebellar: 27.5 years, 95% CI 26.9–28.1) (Fig. 2g,h). The ventricular volume remained stable during early life before expanding in later decades (Fig. 2i). Global cortical surface-based metrics demonstrated progressive maturation from thickness peaking in early childhood (4.2 years, 95% CI 4.0–4.4), to folding peaking in middle childhood (7.5 years, 95% CI 7.0–8.0) and surface area peaking in early adolescence (12.4 years, 95% CI 11.6–13.1) (Supplementary Fig. 1).

**Figure 2.**
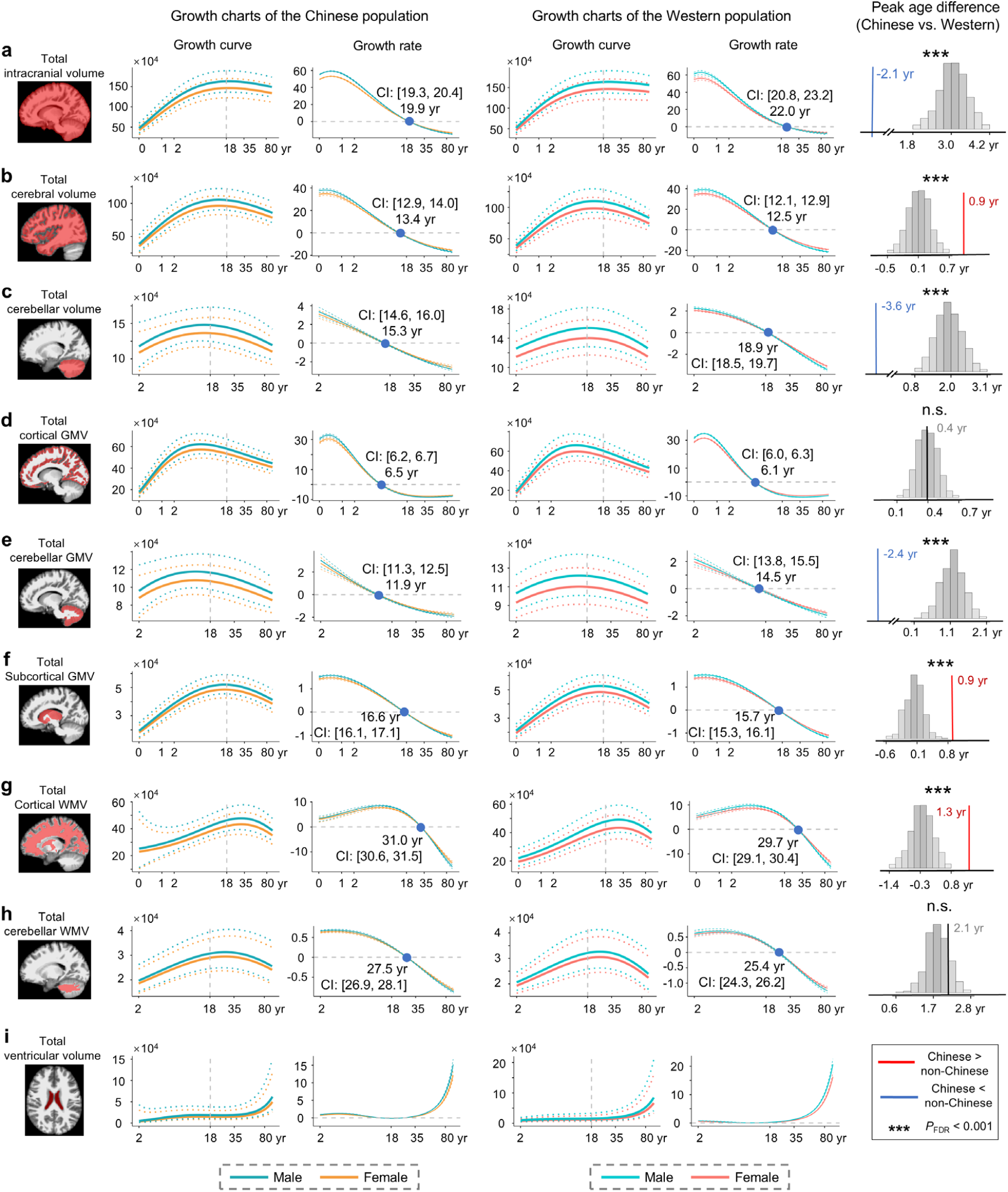
Population-specific growth patterns of global brain phenotypes. Growth patterns of the total intracranial volume (**a**), cerebral volume (**b**), cerebellar volume (**c**), cortical GMV (**d**), cerebellar GMV (**e**), subcortical GMV (**f**), cerebral WMV (**g**), cerebellar WMV (**h**), and ventricular volume (**i**). The panels showed the normative growth curves and growth rates across the lifespan for the Chinese and Western populations. In the growth curve plots, solid lines represent the median (50th percentile), and dotted lines indicate the 5th and 95th percentiles. Growth rates are calculated from the first derivative of the median curves, and 95% confidence intervals (dotted lines) are estimated via 1,000 bootstrap samples (see the Methods for details). Data distributions of these global phenotypes are shown in Supplementary Fig. 2a. The right panels depict differences in the peak age between Chinese and Western models. To quantify differences in the peak age, we performed permutation testing (1,000 iterations) on the pooled data from all Chinese and Western participants, generating a null distribution of peak age differences for each phenotype (see the Methods). GMV, grey matter volume; WMV, white matter volume; CI, confidence intervals; yr, year.

While the Western models also exhibited nonlinear growth trajectories (Fig. 2), permutation analyses (n = 1,000 iterations) revealed significant cross-population variations in maturation peak ages. Specifically, the Chinese population exhibited prolonged cerebral maturation (0.9–1.7 years later, all *P*_*FDR*_ < 0.001) compared with the Western population, involving the total cerebral volume (Fig. 2b), the total subcortical GMV (Fig. 2f), the total cortical WMV (Fig. 2g), and three surface-based global measures (cortical thickness, folding index, and surface area, Supplementary Fig. 1). Conversely, this population demonstrated accelerated maturation (all *P*_*FDR*_ < 0.001) in the total intracranial volume (2.1 years earlier, Fig. 2a), the total cerebellar volume (3.6 years earlier, Fig. 2c), and the total cerebellar GMV (2.4 years earlier, Fig. 2e).

### Population-specific normative growth of subcortical regional phenotypes

Our analysis of Chinese subcortical volumetric development revealed regionally distinct maturation timelines following inverted U-shaped trajectories (Fig. 3 for the left hemisphere). Specifically, the neostriatum (caudate and putamen) exhibited earliest peak maturation, with the caudate peaking in late childhood (9.3 years, 95% CI 8.9–9.5) and the putamen in early adolescence (12.8 years, 95% CI 12.2–13.4), followed by the paleostriatum (pallidum) peaking in early adulthood (21.6 years, 95% CI 20.7–22.6) (Fig. 3a-c). The thalamus and limbic structures reached peak maturation in early adulthood (thalamus: 19.4 years, 95% CI 18.8–20.0; amygdala: 21.7 years, 95% CI 21.2–22.3; hippocampus: 22.3 years, 95% CI 21.8–23.0; Fig. 3d-f). The subcortical maturation patterns of the right hemisphere (Supplementary Fig. 3) were highly similar to those observed in the left hemisphere.

**Figure 3.**
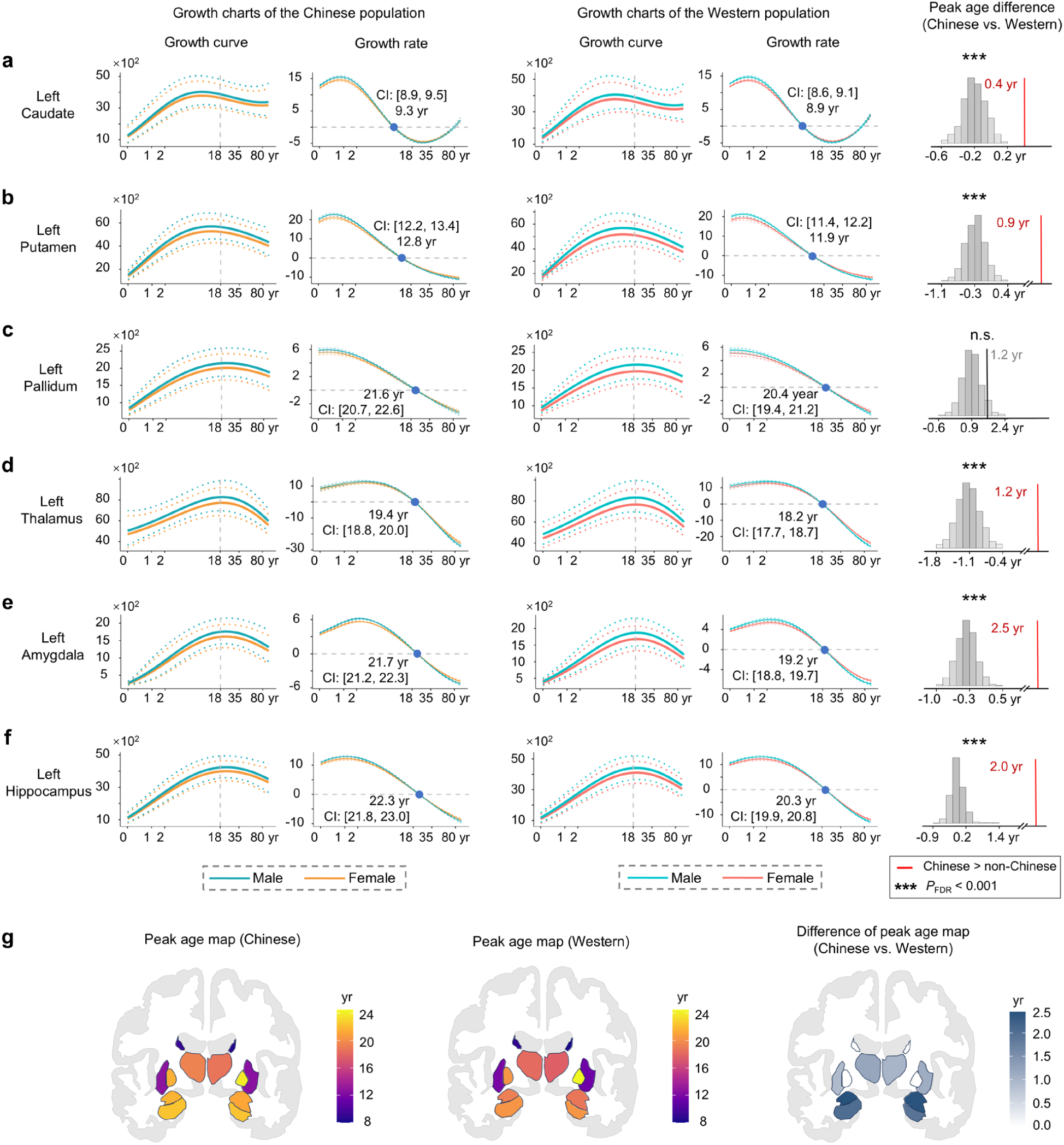
Population-specific growth patterns of subcortical regional phenotypes. **a**, Growth patterns for the GMV of the caudate (**a**), putamen (**b**), pallidum (**c**), thalamus (**d**), amygdala (**e**), and hippocampus (**f**) in the left hemisphere. The panels show the normative growth curves and growth rates across the lifespan for the Chinese and Western populations. In the growth curve plots, solid lines represent the median (50th percentile), and dotted lines indicate the 5th and 95th percentiles. Growth rates are calculated from the first derivative of the median curves, and 95% confidence intervals (dotted lines) are estimated via 1,000 bootstrap samples (see the Methods for details). Data distributions of these subcortical phenotypes are shown in Supplementary Fig. 2b. The right panels depict differences in the peak age between Chinese and Western growth trajectories. To quantify differences in the peak age, we performed permutation testing (1,000 iterations) on the pooled data from all Chinese and Western participants, generating a null distribution of peak age differences for each phenotype (see the Methods for details). **g**, The peak age map of Chinese and Western populations, and the peak age difference map between these two populations. Corresponding results for the right hemisphere are shown in Supplementary Fig. 3. GMV, grey matter volume; CI, confidence intervals; yr, year.

These nonlinear patterns were also exhibited in the Western models (Fig. 3). However, permutation testing revealed significantly prolonged peak maturation in several subcortical structures (caudate, putamen, thalamus, amygdala, and hippocampus) (left hemisphere: 0.4–2.5 years; right hemisphere: 0.3–2.5 years) in the Chinese population compared with the Western population (*P*_FDR_ < 0.001, Fig. 3g and Supplementary Fig. 3).

### Population-specific normative growth of cortical regional phenotypes

We further charted Chinese lifespan cortical development across 68 anatomically defined regions ^22^, revealing three important findings (Fig. 4 and Supplementary Fig. 4). First, we identified an across-feature neurodevelopmental sequence: initial maturation of the cortical thickness, followed by the GMV and gyrification, with the surface area maturing last, paralleling the total phenotypic developmental patterns. Second, regional maturation generally followed a hierarchical developmental sequence, with primary cortices reaching their peak ages earlier than paralimbic association cortices (limbic and insular areas). Third, feature-specific analyses demonstrated strong intercorrelations of peak age maps among the cortical GMV, cortical thickness, and surface area (all r > 0.64, *P* < 0.001), whereas gyrification exhibited selective coupling with the surface area only (r = 0.43, *P* < 0.001) (Supplementary Fig. 5a).

**Figure 4.**
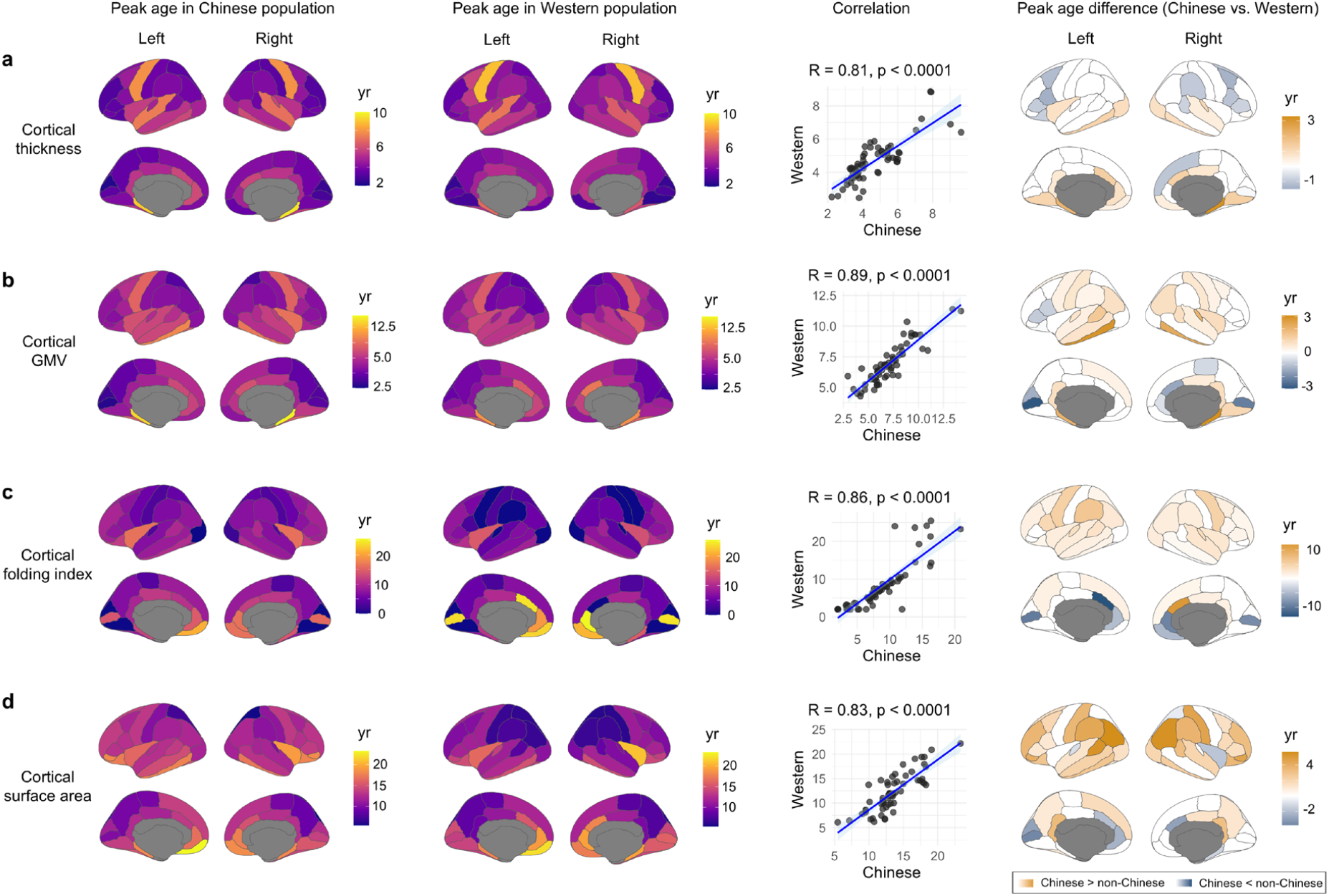
Population-specific growth patterns of cortical regional phenotypes. From top to bottom, the panels correspond to the regional cortical thickness (**a**), GMV (**b**), folding index (**c**), and surface area (**d**). This figure displays: (1) peak maturation ages across 68 cortical regions in Chinese- and Western-specific normative growth curves and their spatial correlations, and (2) regional differences in peak maturation timing between Chinese and Western populations. GMV, grey matter volume.

These patterns were remarkably conserved across populations (Fig. 4 and Supplementary Fig. 5b). However, permutation testing (n = 1,000) revealed population-specific differences: the Chinese population presented prolonged peak maturation in higher-order association cortices (lateral temporal/frontal/parietal cortices, insula, and posterior cingulate) but accelerated development in the primary visual cortex across most metrics. Compared with the Western population, feature-specific exceptions emerged: (1) in the inferior frontal gyrus, the Chinese population presented earlier peaks in the cortical volume and thickness but prolonged surface area maturation, and (2) in the anterior cingulate cortex, earlier peaks in the surface area and gyrification but prolonged thickness maturation were observed in the Chinese population.

### Enhanced phenotypic predictive accuracy using Chinese-specific brain growth models

To evaluate the out-of-sample phenotypic predictive accuracy of the Chinese and Western models for Chinese participants, we employed an independent testing set of healthy individuals from the Chinese case–control datasets (N = 2,591, aged 6–95 years, 26 sites) (Fig. 5a, b). Given that these data were used to train the original Chinese normative growth models, we retrained the models to avoid data leakage. Specifically, we randomly split these Chinese healthy individuals into training and testing subsets (N_train_ = 1,223, N_test_ = 1,368), stratified by age, sex, and site. The Chinese growth chart models were retrained using 41,669 samples for the Chinese population (43,037 - N_test_). For comparison, we also retrained the Western model using 57,562 samples (56,339 + N_train_). The held-out testing subset (N_test_), excluded from model training, was used to assess the predictive performance.

**Figure 5.**
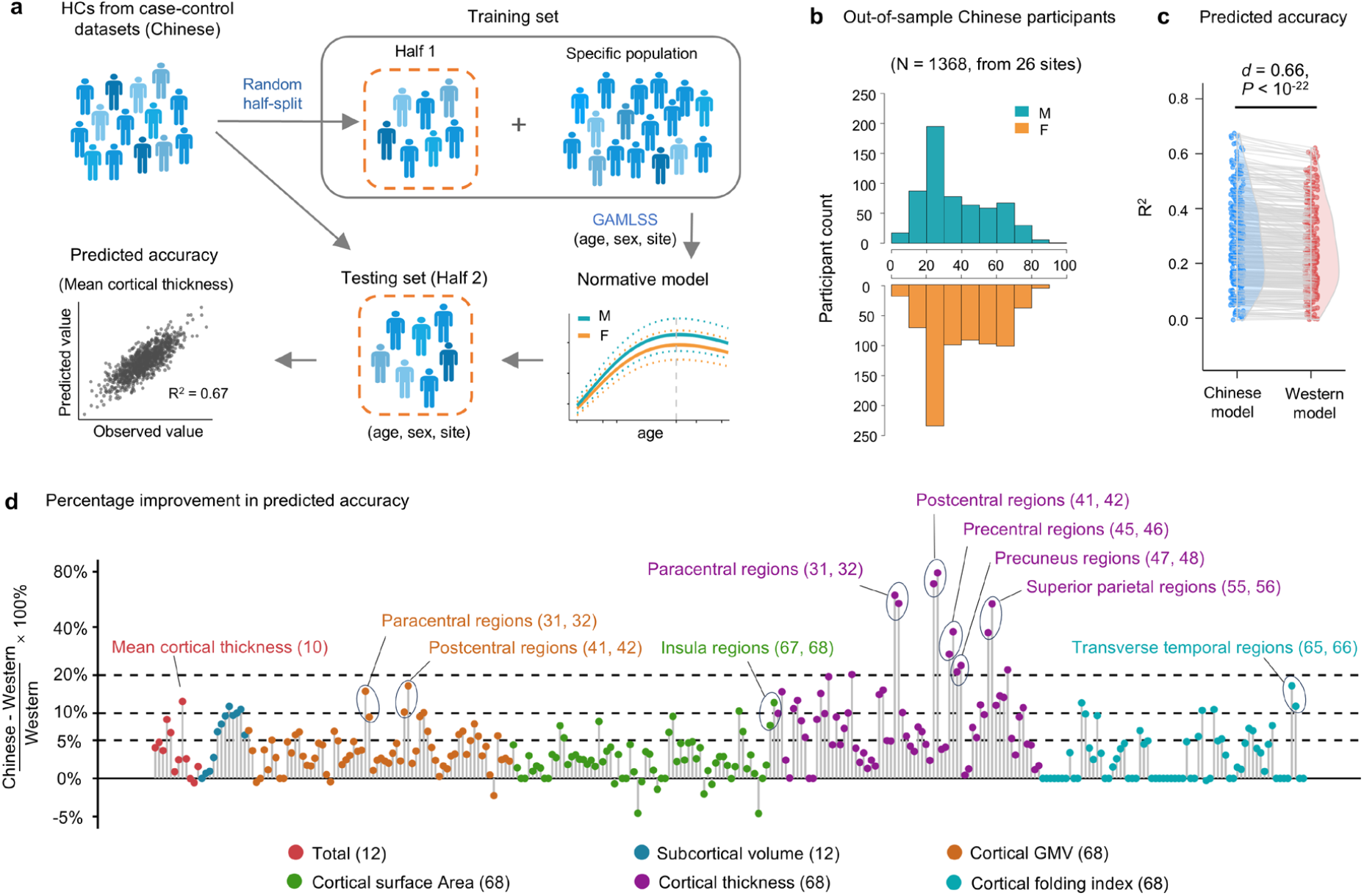
Evaluation of the out-of-sample predictive phenotypic accuracy of Chinese healthy individuals across normative models. **a**, Framework for assessing the predictive accuracy (R^2^) via independent Chinese testing data. **b**, Age and sex distributions of the out-of-sample Chinese participants (testing set). **c**, Comparisons of the predictive accuracy between the Chinese and Western models across all 296 structural brain phenotypes. **d**, Percent improvement in the predictive accuracy of Chinese model than Western model. To avoid spurious values resulting from near-zero R^2^ values in certain phenotypes, improvement rates were computed only for phenotypes with R^2^ > 0.1. The numbers of phenotypes with R^2^ ≥ 0.1 were 253 and 249 in the Chinese and Western models, respectively. The region numbers shown after each brain phenotype correspond to those in Fig. 1e. GMV, grey matter volume.

We quantified the predictive accuracy (R^2^) for each phenotype across the Chinese and Western models, using age, sex, and site as input features for each individual. Overall, the Chinese model demonstrated significantly higher predictive accuracy than the Western model (Cohen’s d = 0.66, mean directional error = 0.84, *P*_FDR_ < 10^−24^) (Fig. 5c). Notably, the Chinese model exhibited at least a 10% improvement in the predictive accuracy over the Western model for 41 phenotypes and at least a 5% improvement for 94 phenotypes (Fig. 5d). The largest improvements (> 20%) occurred in cortical thickness measures of the bilateral paracentral, postcentral, precentral, precuneus, and superior parietal regions (Fig. 5d).

### Reduced misestimation of deviation scores using Chinese-specific brain growth models

Recent studies have highlighted the utility of brain charts in addressing inherent heterogeneity of clinical cohorts by enabling individual-level statistical inference ^1, 3, 6, 7^. These models quantify deviations of individual brain phenotypes from normative expectations, providing insights into neurotypical or atypical variations. Here, we sought to determine whether the Western models would misestimate deviation scores in Chinese clinical cohorts. To do this, we analysed patient data from Chinese case-control datasets encompassing four brain disorders: Alzheimer’s disease (AD; N = 399, aged 43–89 years), mild cognitive impairment (MCI; N = 283, aged 42–88 years), schizophrenia (SCZ; N = 691, aged 11–67 years), and major depressive disorder (MDD; N = 1,492, aged 11–93 years). We computed age- and sex-adjusted deviation *z*-scores for 296 structural brain phenotypes using the abovementioned retrained Chinese and Western models.

In AD patients, the Chinese model captured expected patterns of brain atrophy, including positive deviations in ventricular volume and negative deviations in the hippocampal and amygdala volumes, as well as reduced cortical GMV and thickness, primarily in regions of the precuneus, posterior cingulate, inferior parietal, and temporal cortex (Fig. 6a, b). These findings were highly compatible with previous case–control reports of structural atrophy in AD ^23-25^.

**Figure 6.**
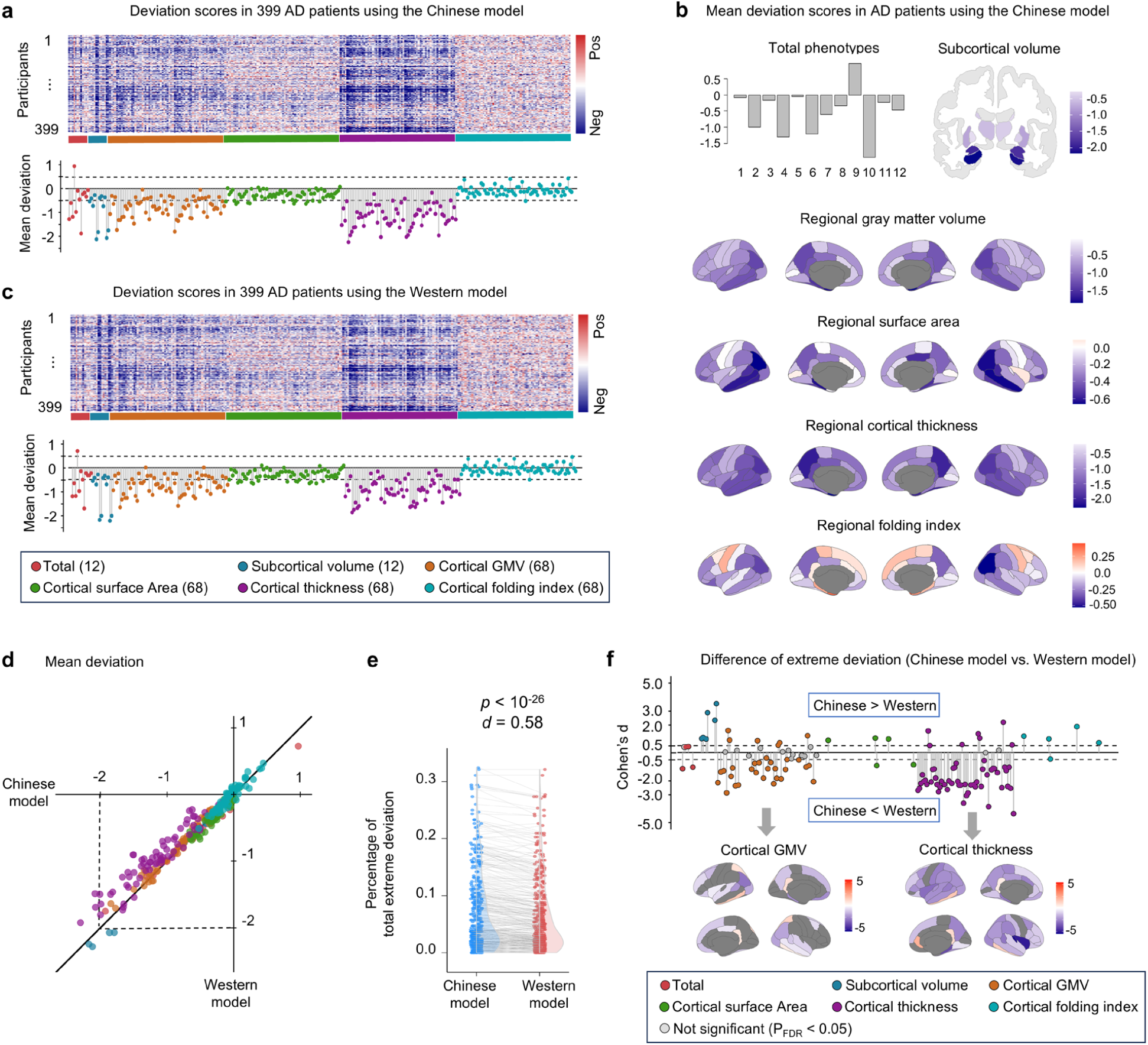
Misestimation of deviation scores in AD by the Western model relative to the Chinese model. **a**, Top panel: Individual-level deviation z scores across 296 phenotypes for 399 AD patients, estimated using the Chinese normative model. Bottom panel: Mean deviation z scores for each phenotype across all AD patients. **b**, Visualization of the mean deviation scores for global, subcortical, and cortical regional phenotypes. The region numbers for the global brain phenotypes correspond to those shown in Fig. 1e. **c**, Corresponding results derived from the Western normative model. **d**, Scatter plot comparing the mean deviation scores between the Chinese and Western models across all phenotypes. **e**, The Chinese model identified a significantly greater proportion of extreme deviations (|z| > 2.6) than did the Western model across AD patients. **f**, For phenotypes with ≥ 20 patients exhibiting extreme deviations in either model, we assessed whether the Western model systematically over- or underestimated the deviation scores. For the cortical GMV and cortical thickness, all group-level mean deviations were negative; thus, negative Cohen’s d values indicate underestimation of the extreme deviation magnitude, whereas positive values reflect overestimation. GMV, grey matter volume.

Similar abnormal patterns were also observed using the Western models (Fig. 6c), whereas the mean deviation extents across all patients tended to be smaller as compared to those obtained using the Chinese model (Fig. 6d). Given the clinical importance of extreme deviations (|z| > 2.6) for identifying neuroanatomical abnormalities ^26, 27^, we next quantified the prevalence of extreme deviations per patient across models. The Chinese models identified significantly more extreme deviations than the Western models (Cohen’s d = 0.58, *P*_FDR_ < 10^−26^; Fig. 6e). To further characterize systematic biases in the Western models, we examined whether specific phenotypes exhibited significant under- or overestimation of extreme deviations. We identified 82 phenotypes that were significantly underestimated and 28 phenotypes that were significantly overestimated in the Western model, with underestimation most prominently affecting measures of the cortical thickness and cortical GMV (Fig. 6f). Split-half replication confirmed the robustness of these results for AD patients (Supplementary Fig. 6). For MCI patients, misestimation patterns similar to those for AD patients were observed, albeit with slightly reduced effect sizes (Supplementary Fig. 7). For both SCZ patients and MDD patients, the Chinese models also identified significantly more extreme deviations than the Western models (SCZ: Cohen’s d = 0.14, *P*_FDR_ = 0.0002; MDD: Cohen’s d = 0.06, *P*_FDR_ = 0.02) (Supplementary Figs. 8-9). Together, these results collectively demonstrate that Chinese-specific normative models provide a more accurate and sensitive characterization of brain structural variations for patients from Chinese cohorts.

### Sensitivity analyses

We validated the lifespan growth trajectories in Chinese and Western populations and their peak age differences for all phenotypes (for details, see the Methods). (*i*) To examine the potential effects of data samples, a bootstrap resampling analysis was performed (1,000 times). (*ii*) To evaluate whether cross-population differences were influenced by unequal sample sizes across age groups, age-stratified balanced resampling was employed to ensure the uniformity of participant numbers in two populations (N_Chinese_ = N_Western_ = 20,770, resampling 1,000 times). (*iii*) To examine the potential effects of specific imaging sites, leave-one-site-out (LOSO) analysis was performed. (*iv*) To validate the reproducibility of our results, a split-half approach was performed. (*v*) To determine whether image quality affected the growth trajectories, we repeated the GAMLSS modelling with Euler number included as an additional covariate. (*vi*) To control for potential ethnic composition effects, we repeated the analysis in strictly defined ethnic subsamples, using only self-identified Han Chinese (N = 20,104) and White individuals (N = 44,915).

For the lifespan brain trajectories, Pearson correlation coefficients were computed between the sensitivity-derived curves and the corresponding main population-specific models, with a sampling interval of 0.01 years. All validation strategies yielded growth trajectories that were highly consistent with the main models (r = 0.90–1.00, Supplementary Table 2). For peak age differences, we assessed the similarity between each validation analysis and the main result across all phenotypes. Specifically, for (*i*) and (*ii*), we computed 1,000,000 pairwise peak age differences by matching 1,000 Chinese and 1,000 Western models; for (*iii*), a total of 66,816 comparisons were generated from all 384 Chinese and 174 Western LOSO models; and for (*iv*), (*v*) and (*vi*), a single comparison of peak age differences was made per strategy. All five analyses reproduced the overall pattern of cross-population peak age differences, with median correlation coefficients of 0.96, 0.94, 0.99, 0.84, 0.87, and 0.66 for strategies (*i*) through (*vi*) (all *P* < 10^−21^), respectively (Supplementary Fig. 10).

## Discussion

The C-LBM Consortium established the first population-specific brain growth charts for Chinese individuals through harmonized analysis of 43,037 participants across 384 sites. Our models reveal precisely timed neurodevelopmental trajectories for 296 structural phenotypes, revealing previously unrecognized population-specific patterns. Crucially, these Chinese-specific models outperform Western-centric models in terms of both the phenotypic predictive accuracy and pathological detection sensitivity for Chinese populations. These findings decisively challenge the presumption of universal neurodevelopmental standards and demonstrate that Western-centric charts systematically misestimate brain development in non-Western populations. By providing validated ethnic-specific references, we address a critical gap in global neuroscience while establishing a framework for developing representative brain atlases. This work not only advances equitable precision medicine but also calls for a paradigm shift towards population-inclusive neuroimaging standards worldwide.

Mapping normative neurodevelopmental trajectories is crucial for pinpointing critical windows of brain plasticity and disorder vulnerability. Several previous large-scale studies (n > 10000) investigated the lifespan trajectories of brain structural phenotypes. For example, Bethlehem et al. ^1^ provided a foundational delineation of the growth milestones for global and regional cerebral volumes, along with global cortical thickness and surface area. Other studies described the developmental trajectories of subcortical structures ^2, 18, 28^, the cerebellum ^18^, and regional cortical thickness ^2, 29^; however, they did not identify the specific growth milestones. Our work comprehensively characterizes the normative developmental trajectories and maturation timelines of 296 structural brain phenotypes. While our findings on global brain volume measurements are consistent with prior reports ^1, 2^, we provide, for the first time, growth milestone patterns for cerebellar structures, subcortical nuclei, and regional cortical indices, including thickness, surface area, and cortical folding. The cerebellar volume peaks in adolescence, with grey matter maturing earlier than white matter. This temporal pattern mirrors that of the cerebrum, suggesting comparable principles of neurodevelopmental timing across these two major brain structures. Subcortical nuclei show marked heterogeneity, with neostriatal structures (e.g., the caudate and putamen) reaching their peak volume earlier than phylogenetically older regions, such as the globus pallidus, thalamus, and hippocampus, aligning with functional MRI evidence of staggered subcortical circuit reorganization ^4, 30^. Cortical regional development follows a hierarchical spatiotemporal gradient from sensorimotor to paralimbic cortices, showing distinct trajectories across morphological features. Together, this detailed mapping offers a more granular understanding of brain maturation and sets the stage for future clinical applications of individualized neurodevelopmental assessment.

Comparative neurodevelopmental analyses of Chinese and Western populations reveal fundamentally conserved trajectories but distinct spatiotemporal maturation timelines. Systematic differences included (i) delayed cerebral and subcortical maturation and accelerated cerebellar development in Chinese individuals and (ii) divergent regional cortical maturation featuring later development of higher-order association cortices but earlier development of the primary visual cortex. These findings substantially extend prior reports of neuroanatomical differences between Chinese and Caucasian populations ^8, 11, 12, 31^ through comprehensive phenotyping and milestone comparisons. These population variations likely arise from complex gene–environment–cultural interactions. Genomic analyses reveal key insights into population-specific brain development. A genome-wide meta-analysis demonstrated gene-specific regulation of nonlinear brain volume trajectories ^18^, whereas population genetics studies revealed significant variations in single nucleotide polymorphism allele frequencies, linkage disequilibrium patterns, and polygenic risk architectures ^32^. The newly developed Chinese pangenome^33^ provides a critical representation of Asian genomic diversity. Furthermore, a neuroimaging genome-wide association study of 3,414 phenotypes identified 38 Chinese-specific genetic associations ^15^ , highlighting ancestry-dependent influences on brain structure and function. Environmental factors, including gene–environment interactions ^19, 34, 35^, and sociocultural factors ^36^ further shape these neurodevelopmental trajectories. These findings underscore the necessity of population-informed frameworks in neuroscience and the development of precision medicine approaches tailored to diverse global populations.

Ethnic bias in normative predictive algorithms is a well-recognized issue in several research domains. Most normative algorithms have historically been derived from Caucasian participants residing in Western cultural contexts, which limits their generalizability to individuals from other ethnic and sociocultural backgrounds. A straightforward and effective solution to mitigate such bias is to include sufficiently large and representative datasets from diverse populations. Obermeyer *et al*. ^37^ demonstrated that a widely used health-risk algorithm systematically underestimates the illness severity of black patients compared with white patients, largely because of label bias arising from the reliance of the algorithm on healthcare costs. In genomics, polygenic risk scores trained predominantly on individuals of European ancestry fail to reliably generalize to non-European populations ^38, 39^, partly owing to differences in the genetic architecture across ancestries, such as allele frequencies and linkage disequilibrium patterns ^32^. Recent advances in human genetics have begun to address this limitation, driven in part by the growing inclusion of large-scale East Asian cohorts in genomic research ^40^. In this study, we demonstrate similar ethnic bias in applications of normative neuroimaging models. Specifically, the Western-specific lifespan model yielded a significantly lower predictive accuracy than the Chinese-specific model when applied to healthy Chinese individuals (aged 0–100 years). Moreover, a comparison of deviation scores revealed that the Western model systematically misestimated the extent of structural brain deviations in Chinese patients. These findings highlight a critical and underrecognized source of bias in neurodevelopmental modelling and call attention to the need for population-representative brain charts to ensure valid and precise clinical evaluations in diverse groups.

Some challenges require further consideration. (1) The fetal stage is a critical period in the full trajectory of human brain development but was not included in this study, mainly due to the limited availability of fetal neuroimaging data, especially the lack of comparable datasets from both Chinese and Western populations. The current C-LBM Consortium represents Phase I of large-scale data collection; subsequent phases will incorporate high-quality fetal imaging data to fill this important gap. (2) This study focused exclusively on Chinese and Western populations. Future research should pursue broader international collaborations and integrate neuroimaging data from diverse racial and ethnic groups to ultimately contribute to the development of more globally representative normative brain charts. (3) Although we have made extensive efforts to collect data across Chinese provinces, sample acquisition remains challenging in certain regions with limited scanners. Future work should aim to include more geographically comprehensive datasets to further improve the population representativeness. (4) Most datasets used in the current study lack complementary information, such as information on cognitive performance, socioeconomic status, environmental exposure, and genetics, which limits the ability to investigate how these factors contribute to shaping lifespan growth trajectories. Prospective, multimodal cohort studies such as CHIMGEN ^41^ are needed across the lifespan to enable a deeper understanding of the biological, environmental, and sociocultural determinants of brain development. (5) This study is based on cross-sectional data, which may result in an underestimation of age-related changes ^42^. The incorporation of densely collected longitudinal data across the lifespan is essential to more accurately capture brain growth trajectories. With continued data integration, the population-specific normative models established here will be periodically updated to provide dynamic, evolving resources.

## Methods

### Chinese lifespan neuroimaging datasets

To delineate the population-specific normative growth chart of the human brain for the Chinese population, we aggregated available structural MRI datasets from across China. Following rigorous quality control, a total of 43,037 healthy participants (24,425 females) from 384 scanner sites across 29 provinces, aged 0 to 100 years, were included in the final analysis (Fig. 1a, c). For individuals with multiple test-retest scans or longitudinal scans, only one scan was used to construct the normative brain growth charts. Among the participants who provided self-reported ethnic information, 98.5% were identified as Han people. Written informed consent was obtained from all participants and/or their legal guardians, and the recruitment procedures were approved by the local ethics committees for each dataset. The detailed demographics of the datasets are presented in Supplementary Table 1.

### Western lifespan neuroimaging datasets

To compare the brain growth patterns between Chinese and Western populations, we also aggregated a large dataset of structural MRI scans from non-Chinese countries. The data were acquired from 174 scanner sites across eight predominantly Western countries: the United States, the United Kingdom, Canada, Ireland, the Netherlands, Belgium, Switzerland, and Australia. Participants who self-identified as Chinese or Asian (2.5%) were first excluded from the non-Chinese datasets. After quality control, 56,339 healthy participants (28,356 females), aged 0 to 100 years, were retained for analysis. Only one scan per participant was used for constructing the normative growth charts. Among the participants who provided ethnic information, 87.8% were identified as white. Written informed consent was obtained from all participants and/or their legal guardians, and the recruitment procedures were approved by the local ethics committees for each dataset. The detailed demographics of the datasets are presented in Supplementary Table 1.

### Image quality control process

We applied a unified four-stage quality control (QC) pipeline ^4^ that combined automated algorithms with expert visual inspection to evaluate the structural MR images across all 125,884 scans. This standardized framework was designed to systematically identify and exclude scans with structural artefacts, thereby improving the data consistency and reliability. A comprehensive description of the QC procedures is available in our previous study ^4^ and at https://github.com/sunlianglong/BrainChart-FC-Lifespan/blob/main/QC/README.md.

### Structural MRI data processing

Ideally, a structural processing pipeline fully unified across all age groups would maximize the methodological consistency. However, the pronounced anatomical differences across the human lifespan—particularly during the perinatal and early infancy stages—render such uniformity impractical. Thus, age-specific processing strategies were employed for early developmental data, while harmonized pipelines were maintained across the remaining lifespan, consistent with our previous work ^4^.

Specifically, for individuals aged two years and older, all structural MRI data underwent uniform preprocessing with FreeSurfer v6.0.0. FreeSurfer software was integrated within the containerized HCP structural preprocessing pipeline (v4.4.0-rc-MOD-e7a6af9) ^43^ as implemented on the QuNex platform (v0.93.2) ^44^. For all datasets except ABCD and UKB, we applied the HCP-recommended PreFreeSurfer pipeline prior to FreeSurfer processing. This stage focused on normalization of anatomical images, which involved (1) brain extraction, (2) denoising, and (3) bias field correction of T1-weighted and, when available, T2-weighted scans. FreeSurfer processing was then conducted on the normalized images to generate cortical surfaces, which mainly included (4) anatomical tissue segmentation, (5) construction of pial, white, and mid-thickness surfaces, (6) surface topological correction, and (7) projection of the surface onto a sphere for surface registration. For the high-quality structural images from the ABCD and UKB datasets, we used the officially released data and performed processing procedure with FreeSurfer v6.0.0 ^45, 46^.

For participants aged 0–2 years, we employed the iBEAT v2.0 pipeline ^47^, which is optimized for early-age neuroimaging data and has demonstrated superior performance in tissue segmentation and cortical reconstruction for infants compared with alternative approaches ^47^. This pipeline follows principles similar to those of the HCP-FreeSurfer approach and mainly includes (1) inhomogeneity correction of T1w/T2w images, (2) skull stripping, (3) cerebellum removal (for participants with incomplete cerebellum removal, frame-by-frame manual correction was performed), (4) tissue segmentation, (5) cortical surface reconstruction, (6) topological correction of the white matter surface, and (7) final reconstruction of the inner and outer cortical surfaces. For neonates from the dHCP study, we used data processed with the officially recommended dHCP pipeline ^48^, which was specifically designed to account for the substantial differences between neonatal and adult MRI data. This pipeline, which is similar in structure to those described above, mainly includes (1) bias correction and (2) brain extraction, which are performed on motion-corrected, reconstructed T2w images, (3) tissue segmentation, (4) cortical reconstruction of the white matter surface, (5) surface topology correction, (6) generation of pial and mid-thickness surfaces, and (7) spherical projection for surface registration.

The individual cortical surfaces from participants aged 0–2 years were aligned with the adult fsaverage standard space using a three-step registration method ^4^. Briefly, for neonates aged 40 to 44 postmenstrual weeks in the dHCP study, the following steps were implemented. (1) Individual surfaces were registered to their respective postmenstrual week templates ^49^. (2) Templates for 41-44 postmenstrual weeks were registered to the 40-week template. (3) The 40-week template was then registered to the fsaverage surface template. For infants aged 1–24 months, the following steps were taken: (1) Individual surfaces were registered to their corresponding monthly age templates ^50^. (2) All monthly templates were registered to the 12-month template. (3)The 12-month template was then registered to the fsaverage surface template.

### Structural brain phenotypes

#### (i) Global brain phenotypes

We systematically quantified whole-brain global structural measures, including the total intracranial volume, total cerebral volume, total cerebellar volume, cortical GMV, subcortical GMV, cerebellar GMV, cortical WMV, cerebellar WMV, and ventricular volume. The surface-based global measures included the mean cortical thickness, total cortical surface area, and total folding index. The intracranial volume was defined as the volume enclosed within the inner skull surface. In accordance with a previous study ^1^, the total cerebral volume was a proxy as the sum of cortical GM and WM volumes, which is a measure highly correlated with the directly segmented cerebral volume (r = 0.99). Because the iBEAT pipeline does not provide accurate segmentation of ventricles and the cerebellum in infants aged 0–2 years, ventricular and cerebellar metrics were modelled only from 2–100 years of age.

#### (ii) Subcortical regional phenotypes

On the basis of FreeSurfer segmentation, subcortical GM volumes (bilateral caudate, putamen, pallidum, thalamus, amygdala, and hippocampus) were extracted for participants aged 2–100 years. For infants aged 0–2 years, we registered individual structural images to a 12-month template using age-specific registration strategies ^4^, and mapped a standard subcortical atlas to derive individual volumetric measures.

#### (iii) Cortical regional phenotypes

After the reconstruction of the cortical surfaces, the vertex-wise cortical thickness, GM volume, surface area, and folding index were computed for each participant. Individual-to-template surface mappings, as established during data preprocessing (see the “**Structural MRI data processing**” section), were used to project the standard Desikan– Killiany atlas onto each participant’s surface, enabling the extraction of region-wise surface-based measures across 68 cortical regions. Owing to notoriously poor signal quality or missing values in neonatal MRI scans, the entorhinal region, temporal pole region, and frontal pole region presented exceptionally high between-subject variability and outlier-level peak ages ^1^. Thus, these regions were excluded from the set of regional volumetric milestones in the lifespan brain chart proposed by Bethlehem et al. ^1^. Consistent with those findings, we observed similarly unreliable estimates in these regions in our dataset. To maintain methodological alignment and ensure interpretability, we also excluded these three regions from the milestone analyses.

### Modelling the lifespan growth curves

To estimate the normative growth patterns of various structural phenotypes in healthy Chinese individuals, we applied generalized additive models for location, scale and shape (GAMLSS) ^51, 52^ using the *gamlss* package (version 5.4-3) in R 4.2.0. GAMLSS, recommended by the World Health Organization for normative modelling, serves as a robust statistical tool for delineating complex, nonlinear growth patterns, with broad applications in neurodevelopmental studies ^1, 4, 11^.

#### (i) GAMLSS framework

Recent work by Bethlehem et al ^1^ introduced a robust GAMLSS framework and demonstrated that the generalized gamma (GG) distribution is the most suitable for brain structural phenotypic data. The GG distribution is characterized by three parameters: the median (*µ*), coefficient of variation (*σ*), and skewness (*ν*). On the basis of this data distribution, we determined the best-fitting parameters for each structural phenotype. Specifically, to flexibly model age-related changes, we employed the fractional polynomial function as the smoothing term within the GAMLSS framework, which allowed the simultaneous fitting and selection of nonlinear terms during model estimation. To ensure optimal model fit, we systematically explored alternative model configurations by varying the number of polynomials (ranging from 1 to 3) used to parameterize the μ and *σ* components of the GG distribution, and additionally selected whether to include random effects for site in these parameters. Following Bethlehem et al ^1^, polynomials for *ν* were not evaluated, as modelling *ν* with fractional polynomials led to instability in model selection (e.g., failure to converge to an optimal parameter), and there was no a priori reason to assume age- or random-effect-dependent changes in skewness. For model estimation, we adopted the default convergence criterion of a log-likelihood = 0.001 between iterations and set the maximum number of iterations at 200. The Bayesian information criterion (BIC) was used to evaluate the fit of all the converged models, and the model with the lowest BIC was selected as the optimal model. No convergence failures were observed across any structural phenotypes. As an illustrative example, we present the final model specification for the total intracranial volume.

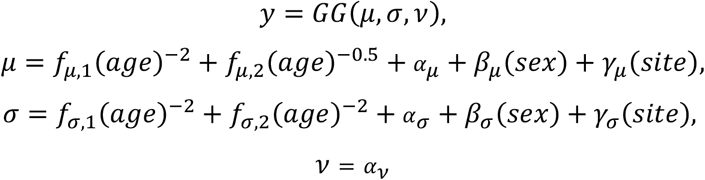

The *f* terms describe the fixed effects of age, modelled using fractional polynomials, with the number of terms indicating the polynomial order. In accordance with Bethlehem et al. ^1^, age was measured in days and log-transformed to emphasize the rapid developmental changes during early life. The *α* terms represent the fixed intercepts, the *β* terms represent the fixed effects of sex, and the *γ* terms represent site-level random effects. For the skewness parameter (*ν*), only a fixed intercept was included. Using the above GAMLSS framework, we also estimated Western normative models for each phenotype on basis of the Western population datasets.

### Differences in the peak age between the Chinese and Western growth curves

To assess whether the peak ages of the Chinese-specific and Western-specific lifespan growth curves significantly differed, we performed a permutation test. Specifically, all Chinese (N = 44,969) and Western (N = 56,339) participants were combined into a single pool. Age was stratified into 26 intervals (0–3 months, 3–6 months, 6–12 months, 1–2 years, 2–4 years, 4–6 years, 6–8 years, 8–10 years, 10–15 years, 15–20 years, 20–25 years, 25–30 years, 30–35 years, 35–40 years, 40–45 years, 45–50 years, 50–55 years, 55–60 years, 60–65 years, 65–70 years, 70–75 years, 75–80 years, 80–85 years, 85–90 years, 90–95 years, and 95–100 years). Stratified random sampling was performed on the basis of age and sex, with 44,969 participants assigned to the “Chinese” group and the remainder assigned to the “Western” group. Normative growth curves were then separately re-estimated for these two pseudo-groups, and the peak age difference between the two curves was calculated. This permutation procedure was repeated 1,000 times, generating a null distribution of the peak age differences for each phenotype. The empirical p value for each phenotype was determined on basis of the position of the observed peak age difference relative to the null distribution. Multiple comparisons across all phenotypes were corrected using a false discovery rate (FDR) threshold of 0.05, with phenotypes surviving correction considered to show significant peak age differences.

### Sensitivity analysis of normative growth models

The Chinese-specific and Western-specific normative growth trajectories, as well as their peak age differences, were validated using multiple sensitivity analyses. These analyses addressed key methodological concerns, including model stability, validation using age-matched subsamples with equal sample sizes across the two populations, potential site-specific effects, replication in independent subsamples, assessment of imaging quality influence, and validation using more strictly defined ethnic subsamples (self-reported Han Chinese and white individuals). For the population-specific growth trajectories, we quantitatively assessed the similarity between these validated growth patterns and the main results by sampling each growth curve at 1,000 points, and computing the Pearson’s correlation coefficients between corresponding curves.

#### (i) Bootstrap resampling analysis

To evaluate the robustness of the lifespan growth curves and estimate confidence intervals, we performed bootstrap resampling separately for the Chinese and Western populations. A total of 1,000 bootstrap samples were generated using replacement sampling ^1^. For each bootstrap sample, normative growth models were refitted for all phenotypes, yielding 1,000 growth trajectories per phenotype. We first assessed the similarity between each bootstrap-derived trajectory and the original model to evaluate the within-population model stability. To assess the robustness of cross-population differences, we obtained all pairwise differences in the peak age between the 1,000 Chinese and 1,000 Western bootstrap trajectories (1,000 × 1,000 = 1,000,000 comparisons) and calculated the correlation between each pairwise peak age difference and the main result across all phenotypes.

#### (ii) Age-matched samples sizes across two populations

To evaluate whether cross-population differences were influenced by unequal sample sizes across age groups, we conducted a sensitivity analysis using age-stratified balanced resampling. Specifically, for each of the 26 age intervals (e.g., 0–3 months, 3–6 months, …, 95–100 years; see the “**Peak age difference between Chinese and Western growth curves**” section), we matched the numbers of Chinese and Western participants by randomly downsampling the group with more samples to match the size of the smaller group. This procedure was repeated 1,000 times. In each iteration, the total number of participants was fixed at 20,770 per population. For each of the 1,000 resampled datasets, Chinese- and Western-specific normative models were re-estimated for all the phenotypes. To assess the within-population model stability, we computed the similarity between each resampled growth trajectory and the original model. To assess the robustness of cross-population differences, we obtained all pairwise differences in the peak age between the 1,000 Chinese and 1,000 Western growth trajectories (1,000 × 1,000 = 1,000,000 comparisons) and calculated the correlation between each pairwise peak age difference and the main result across all phenotypes.

#### (iii) Leave-one-site-out analysis

To determine whether the lifespan growth curves were influenced by data from specific imaging sites, we conducted leave-one-site-out (LOSO) analyses for each phenotype and each population-specific model. In each iteration, all samples from a single site were excluded, and the GAMLSS was re-estimated on the basis of the remaining data to derive a new growth trajectory. This procedure resulted in 384 LOSO models for the Chinese population and 174 for the Western population. We first evaluated the similarity between each LOSO-derived growth curve and the original model to assess the within-population stability. To test the robustness of cross-population differences, we obtained all pairwise differences in the peak age between the 384 Chinese and 174 Western LOSO trajectories (384 × 174 = 66,816 comparisons) and calculated the correlation between each pairwise peak age difference and the main result across all phenotypes.

#### (iv) Split-half replication analysis

To assess the model replicability in independent datasets, a split-half strategy was used. The participants from each population were randomly split into two subgroups stratified by age, sex and site. Each subgroup comprised approximately 50% of the total sample (Chinese: N_1_ = 21,840, N2 = 21,197; Western: N_1_ = 28,104, N2 = 28,235). Lifespan normative growth patterns were independently estimated within each subgroup for both the Chinese and Western populations. For each phenotype, we assessed the similarity of the growth trajectories and the cross-population peak age differences between each subgroup and the main results, as well as between the two subgroups.

#### (v) Imaging quality–controlled modelling

To assess whether image quality influenced the lifespan growth trajectories, we included the Euler number—a widely used proxy for cortical surface reconstruction quality ^53^—as an additional covariate in the GAMLSS models for each phenotype and population. We then compared the resulting trajectories and peak age differences with those from the original models to evaluate the impact of image quality.

#### (vi) Validation analysis using Han Chinese and white populations

Among the Chinese participants with self-reported ethnicity (N = 20,410), 98.5% identified as Han Chinese. Among the Western participants with self-reported ethnicity (N = 51,156), 87.8% identified as white. Although these groups constituted the vast majority of their respective populations, we conducted a stricter validation analysis to control for potential ethnic composition effects. Specifically, we selected only those participants who explicitly identified as Han Chinese (N = 20,104) or white (N = 44,915) and re-estimated all normative models using these subsamples. For each phenotype, we assessed the similarity of the resulting lifespan growth trajectories and corresponding peak ages to those of the full population models. Notably, because few participants younger than 6 had available ethnicity information in our dataset, all Han Chinese- and White-specific models were fitted using data from individuals aged 6 to 100 years. Accordingly, comparisons with the main models were restricted to this age range (6–100 years). A total of 154 phenotypes with original peak ages occurring after age 6 were retained for the association analyses.

### Predictive accuracy for brain phenotypes of the Chinese and Western growth models

To examine whether the Chinese-specific growth model better captures the brain structural features of Chinese individuals than the Western models, we evaluated the out-of-sample predictive accuracy of each model. We employed an independent testing set of healthy controls from the Chinese case–control datasets (N = 2,591, aged 6–95 years, 26 sites) (Fig. 5b). Since these individuals were part of the original Chinese normative model training set, all models were retrained to avoid data leakage. Specifically, the 2,591 healthy Chinese controls were randomly split into training (N_train_ = 1,223) and testing (N_test_ = 1,368) subsets, which were stratified by age, sex, and site. The retrained models used 41,669 samples for the Chinese population (43,037 − N_test_) and 57,562 samples for the Western population (56,339 + N_train_). The held-out testing subset (N_test_), excluded from model fitting, was used to assess the predictive performance of all three models (Fig. 5a).

For a given phenotype, *y* is the observed value. Each participant’s age, sex, and site information were input into the phenotype-specific normative model to generate predicted values *ŷ*. The predictive accuracy was quantified using the coefficient of determination *R*^*2*^, which was calculated as *R*^*2*^ = 1−SSE/SST, where SSE is the sum of squared errors between the observed and predicted values and SST is the total sum of squares relative to the mean of the observed values. After *R*^*2*^ was calculated for all phenotypes under each model, pairwise comparisons were performed to evaluate the differences in the predictive accuracy between models, including the effect size and statistical significance. We further calculated the percentage improvement in the predictive performance of the Chinese model relative to that of the Western model. To avoid spurious values resulting from near-zero R^2^ values in certain phenotypes, improvement rates were computed only for phenotypes with R^2^ > 0.1. The improvement was computed as 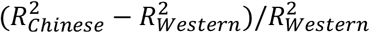.

### Clinical relevance of Chinese-specific normative models for brain disorders

To further assess the clinical utility of the Chinese-specific lifespan growth models, we incorporated quality-controlled structural MRI data for four brain disorders. All quality control and preprocessing procedures were identical to those applied during normative model construction for healthy individuals. The final dataset included 399 patients with Alzheimer’s disease (AD; aged 43–89 years), 283 patients with mild cognitive impairment (MCI; aged 42–88 years), 691 patients with schizophrenia (SCZ; aged 11–67 years), and 1,492 patients with major depressive disorder (MDD; aged 11–93 years).

On the basis of three retrained models described in the “**Predictive accuracy for brain phenotypes of the Chinese and Western growth models**” section, patient data were used as an independent testing set for estimation of individual deviation scores. Specifically, for each patient, quantile scores were first calculated relative to the normative growth curves of each model ^1^. Deviation z scores were then derived using quantile randomized residuals ^54^, which transformed quantiles from the fitted GG distribution into z scores following a standard Gaussian distribution. The normality of the resulting deviation z scores was assessed using a two-tailed Kolmogorov–Smirnov test, confirming that the z scores followed a normal distribution across all phenotypes (P_FDR_ < 0.05). Extreme deviations were defined as z > |2.6| (corresponding to *P* < 0.005), which is consistent with the criteria used in previous studies ^26, 27^. The percentage of total extreme deviation scores across all phenotypes was calculated for each patient.

Given the reduced predictive accuracy of the Western models for healthy Chinese population (Fig. 5), we further investigated whether these models would similarly misestimate the deviation scores in Chinese clinical cohorts. Differences in the proportion of extreme deviations between the Chinese models and the Western models were assessed using two-sample t tests, and the effect sizes were calculated. For each disorder, and for each phenotype with at least 20 patients exhibiting extreme deviations, we compared the deviation scores between models to quantify model-specific biases. To evaluate the robustness of these findings, we performed split-half replication for AD patients. Patients were randomly divided into two subgroups, and all analyses were repeated. Consistent results were observed across subgroups, supporting the stability of the estimates of the Chinese model for clinical populations.

## Data Availability

The MRI dataset listed in Supplementary Table 1 are partially publicly available at the Adolescent Brain Cognitive Development Study (https://nda.nih.gov/), the Autism Brain Imaging Data Exchange Initiative (https://fcon_1000.projects.nitrc.org/indi/abide/), the ADHD200 (https://fcon_1000.projects.nitrc.org/indi/adhd200/), the Alzheimer’s Disease Neuroimaging Initiative (https://adni.loni.usc.edu/), the Age_ility Project (https://www.nitrc.org/projects/age-ility), the Baby Connectome Project (https://nda.nih.gov/), the Brain Genomics Superstruct Project (https://doi.org/10.7910/DVN/25833), the Calgary Preschool MRI Dataset (https://osf.io/axz5r/), the Cambridge Centre for Ageing and Neuroscience Dataset (https://www.cam-can.org/index.php?content=dataset), the Chinese Color Nest Project (https://www.scidb.cn/en/detail?dataSetId=c81f0e90a51b4cfca348ce4da6ca734e), the Chinese Human Connectome Project (https://www.scidb.cn/en/detail?dataSetId=f512d085f3d3452a9b14689e9997ca94), the Cincinnati MR Imaging of Neurodevelopment (https://nda.nih.gov/edit_collection.html?id=2329); the Consortium for Reliability and Reproducibility (https://fcon_1000.projects.nitrc.org/indi/CoRR/html/), the Dallas Lifespan Brain Study (https://fcon_1000.projects.nitrc.org/indi/retro/dlbs.html), the Developing Human Connectome Project (http://www.developingconnectome.org/data-release/second-data-release/), the University of North Carolina Early Brain Development Study (https://www.med.unc.edu/psych/research/psychiatry-department-research-programs/early-brain-development-research/), the FBIRN Phase II Multi-site fMRI programs (http://www.nbirn.net/Resources/Downloads), the Healthy Brain Network (https://fcon_1000.projects.nitrc.org/indi/cmi_healthy_brain_network/MRI_EEG.html), the Human Connectome Project (https://www.humanconnectome.org), the Lifespan Human Connectome Project (https://nda.nih.gov/), the Nathan Kline Institute-Rockland Sample Dataset (https://fcon_1000.projects.nitrc.org/indi/pro/nki.html), the Neuroscience in Psychiatry Network Dataset (https://nspn.org.uk/), the Open Access Series of Imaging Studies (https://sites.wustl.edu/oasisbrains/), the Pediatric Imaging, Neurocognition, and Genetics (PING) Data Repository (http://pingstudy.ucsd.edu/), the Pixar Dataset (https://openfmri.org/dataset/ds000228/), the Philadelphia Neurodevelopmental Cohort (https://www.med.upenn.edu/bbl/philadelphianeurodevelopmentalcohort.html), the Queensland Twin Adolescent Brain (https://openneuro.org/datasets/ds004146/versions/1.0.4), the Strategic Research Program for Brain Sciences MRI Dataset (https://bicr-resource.atr.jp/srpbsopen/), the OpenfMRI database ds000115 (https://openfmri.org/dataset/ds000115/), the SchizConnect Project (including the BrainGluSchi, the COBRE, the MCICShare, and the NMorphCH dataset, https://schizconnect.org/), the Southwest University Adult Lifespan Dataset (http://fcon_1000.projects.nitrc.org/indi/retro/sald.html), the Southwest University Longitudinal Imaging Multimodal Brain Data Repository (http://fcon_1000.projects.nitrc.org/indi/retro/southwestuni_qiu_index.html), and the UK Biobank Brain Imaging Dataset (https://www.ukbiobank.ac.uk/). The dhcpSym surface atlases in aged from 40 to 44 postmenstrual weeks is available at https://brain-development.org/brain-atlases/atlases-from-the-dhcp-project/cortical-surface-template/. The UNC 4D infant cortical surface atlases are available at https://bbm.web.unc.edu/tools/. The fs_LR_32k surface atlas is available at https://balsa.wustl.edu/. The brain charts of the Chinese population are shared online via GitHub (https://github.com/sunlianglong/Population-specific-brain-charts).

## Code Availability

The codes for this manuscript are available on GitHub (https://github.com/sunlianglong/Population-specific-brain-charts). Software packages used in this manuscript include MRIQC v0.15.0 (https://github.com/nipreps/mriqc), QuNex v0.93.2 (https://gitlab.qunex.yale.edu/), HCP pipeline v4.4.0-rc-MOD-e7a6af9 (https://github.com/Washington-University/HCPpipelines/releases), ABCD-HCP pipeline v1 (https://github.com/DCAN-Labs/abcd-hcp-pipeline), dHCP structural pipeline v1 (https://github.com/BioMedIA/dhcp-structural-pipeline), iBEAT pipeline v2.0.0 (https://github.com/iBEAT-V2/iBEAT-V2.0-Docker), MSM v3.0 (https://github.com/ecr05/MSM_HOCR), FreeSurfer v6.0.0 (https://surfer.nmr.mgh.harvard.edu/), FSL v6.0.5 (https://fsl.fmrib.ox.ac.uk/fsl/fslwiki), Connectome Workbench v1.5.0 (https://www.humanconnectome.org/software/connectome-workbench), MATLAB R2018b (https://www.mathworks.com/products/matlab.html), BrainNet Viewer toolbox v 20191031 (https://www.nitrc.org/projects/bnv), R v4.2.0 (https://www.r-project.org), GAMLSS package v5.4-3 (https://www.gamlss.com/),and ggplot2 package v3.4.2 (https://ggplot2.tidyverse.org/). The codes for the GAMSS model are from Bethlehem et al. ^1^ at GitHub (https://github.com/brainchart/Lifespan).

## Acknowledgments

This work was supported by grants from the National Natural Science Foundation of China (82021004, 82327807 and 31830034 to Y. He, T24B2012 to L. Sun, 824B2051 to X. Liang, 82472052 to W. Qin, and 82430063 to C. Yu), the National Key Research and Development Program of China (2018YFC1314300 to C. Yu), the scientific and technological innovation 2030 - the major project of the Brain Science and Brain-Inspired Intelligence Technology (2021ZD0200500 to Q. Dong). We thank Xinian Zuo, Xinlin Zhou, Dairong Cao, Xiangrong Zhang, Tao Wu, and Keith Kendrick for collecting a subset of the data for this study. We thank Liyuan Lin, Chenxuan Pang, Qian Wang, Qian Yu, Maolin Li for their assistance with the visual inspection component of data quality control. We are grateful to the Adolescent Brain Cognitive Development (ABCD) Study, the Autism Brain Imaging Data Exchange (ABIDE) Initiative, the ADHD200 Initiative, the Alzheimer’s Disease Neuroimaging Initiative (ADNI), the Age_ility Project, the Baby Connectome Project (BCP), the Brain Genomics Superstruct Project (BGSP), the Calgary Preschool MRI Dataset, the Cambridge Centre for Ageing and Neuroscience (Cam-CAN) Dataset, the Beijing Cohort in Children Brain Development Project (CBD), the Chinese Color Nest Project (CCNP), the Chinese Human Connectome Project (CHCP), the Cincinnati MR Imaging of Neurodevelopment Dataset (C-MIND), the Consortium for Reliability and Reproducibility (CORR), the Dallas Lifespan Brain Study (DLBS), the developing Human Connectome Project (dHCP), the University of North Carolina Early Brain Development Study (EBDS), the FBIRN Phase II Multi-site fMRI programs, the Healthy Brain Network (HBN), the Human Connectome Project (HCP), the Lifespan Human Connectome Project (HCPA & HCPD), the Nathan Kline Institute-Rockland Sample (NKI-RS) Dataset, the Neuroscience in Psychiatry Network (NSPN) Dataset, the Open Access Series of Imaging Studies (OASIS), the OpenfMRI database ds000115, the Pixar Dataset, the Philadelphia Neurodevelopmental Cohort (PNC), the Queensland Twin Adolescent Brain (QTAB), the Southwest University Adult Lifespan Dataset (SALD), the SchizConnect Project, the Southwest University Longitudinal Imaging Multimodal (SLIM) Brain Data Repository, the UCLA dataset, the UK Biobank (UKB) Brain Imaging Dataset, the Disease Imaging Data Archiving: major depressive disorder (DIDA-MDD) Working Group, the Multi-center Alzheimer Disease Imaging (MCADI) Consortium, the Chinese Imaging Genetics (CHIMGEN) Consortium, and the Chinese Lifespan Brain Mapping (C-LBM) Consortium.

ABCD: data used in the preparation of this article were obtained from the Adolescent Brain Cognitive Development Study (https://abcdstudy.org), held in the NIMH Data Archive (NDA). This is a multisite, longitudinal study designed to recruit more than 10,000 children age 9-10 and follow them over 10 years into early adulthood. The ABCD Study® is supported by the National Institutes of Health and additional federal partners under award numbers U01DA041048, U01DA050989, U01DA051016, U01DA041022, U01DA051018, U01DA051037, U01DA050987, U01DA041174, U01DA041106, U01DA041117, U01DA041028, U01DA041134, U01DA050988, U01DA051039, U01DA041156, U01DA041025, U01DA041120, U01DA051038, U01DA041148, U01DA041093, U01DA041089, U24DA041123, U24DA041147. A full list of supporters is available at https://abcdstudy.org/federal-partners.html. A listing of participating sites and a complete listing of the study investigators can be found at https://abcdstudy.org/consortium_members/. ABCD consortium investigators designed and implemented the study and/or provided data but did not necessarily participate in the analysis or writing of this report. This manuscript reflects the views of the authors and may not reflect the opinions or views of the NIH or ABCD consortium investigators. The ABCD data repository grows and changes over time.

ABIDE I: primary support for the work by Adriana Di Martino was provided by the (NIMH K23MH087770) and the Leon Levy Foundation. Primary support for the work by Michael P. Milham and the INDI team was provided by gifts from Joseph P. Healy and the Stavros Niarchos Foundation to the Child Mind Institute, as well as by an NIMH award to MPM (NIMH R03MH096321).

ABIDE II: primary support for the work by Adriana Di Martino and her team was provided by the National Institute of Mental Health (NIMH 5R21MH107045). Primary support for the work by Michael P. Milham and his team provided by the National Institute of Mental Health (NIMH 5R21MH107045); Nathan S. Kline Institute of Psychiatric Research). Additional Support was provided by gifts from Joseph P. Healey, Phyllis Green and Randolph Cowen to the Child Mind Institute.

ADNI: Data used in preparation of this article were obtained from the Alzheimer’s Disease Neuroimaging Initiative (ADNI) database (adni.loni.usc.edu). As such, the investigators within the ADNI contributed to the design and implementation of ADNI and/or provided data but did not participate in analysis or writing of this report. A complete listing of ADNI investigators can be found at: http://adni.loni.usc.edu/wp-content/uploads/how_to_apply/ADNI_Acknowledgement_List.pdf. Data collection and sharing for this project was funded by the Alzheimer’s Disease Neuroimaging Initiative (ADNI) (National Institutes of Health Grant U01 AG024904) and DOD ADNI (Department of Defense award number W81XWH-12-2-0012). ADNI is funded by the National Institute on Aging, the National Institute of Biomedical Imaging and Bioengineering, and through generous contributions from the following: AbbVie, Alzheimer’s Association; Alzheimer’s Drug Discovery Foundation; Araclon Biotech; BioClinica, Inc.; Biogen; Bristol-Myers Squibb Company; CereSpir, Inc.; Cogstate; Eisai Inc.; Elan Pharmaceuticals, Inc.; Eli Lilly and Company; EuroImmun; F. Hoffmann-La Roche Ltd and its affiliated company Genentech, Inc.; Fujirebio; GE Healthcare; IXICO Ltd.;Janssen Alzheimer Immunotherapy Research & Development, LLC.; Johnson & Johnson Pharmaceutical Research & Development LLC.; Lumosity; Lundbeck; Merck & Co., Inc.; Meso Scale Diagnostics, LLC.; NeuroRx Research; Neurotrack Technologies; Novartis Pharmaceuticals Corporation; Pfizer Inc.; Piramal Imaging; Servier; Takeda Pharmaceutical Company; and Transition Therapeutics. The Canadian Institutes of Health Research is providing funds to support ADNI clinical sites in Canada. Private sector contributions are facilitated by the Foundation for the National Institutes of Health (www.fnih.org). The grantee organization is the Northern California Institute for Research and Education, and the study is coordinated by the Alzheimer’s Therapeutic Research Institute at the University of Southern California. ADNI data are disseminated by the Laboratory for Neuro Imaging at the University of Southern California.

BCP: data used herein is supported by NIH grant (1U01MH110274) and the efforts of the UNC/UMN Baby Connectome Project Consortium.

dHCP: data were provided by the developing Human Connectome Project, KCL-Imperial-Oxford Consortium funded by the European Research Council under the European Union Seventh Framework Programme (FP/2007-2013) / ERC Grant Agreement no. [319456]. We are grateful to the families who generously supported this trial.

HCP: data were provided by the Human Connectome Project, WU-Minn Consortium (Principal Investigators: David Van Essen and Kamil Ugurbil; 1U54MH091657) funded by the 16 NIH Institutes and Centers which support the NIH Blueprint for Neuroscience Research; and by the Mc-Donnell Center for Systems Neuroscience at Washington University.

HCP Lifespan: data used in this publication was supported by the National Institute of Mental Health of the National Institutes of Health under Award Number U01MH109589 and by funds provided by the McDonnell Center for Systems Neuroscience at Washington University in St. Louis. The content is solely the responsibility of the authors and does not necessarily represent the official views of the National Institutes of Health.

NKI-RS: funding for key personnel provided in part by the New York State Office of Mental Health and Research Foundation for Mental Hygiene. Additional project support provided by the NKI Center for Advanced Brain Imaging (CABI), the Brain Research Foundation (Chicago, IL), the Stavros Niarchos Foundation, and NIH grant P50 MH086385-S1.

NSPN: the NSPN study was funded by a Wellcome Trust award to the University of Cambridge and the University College London.

UK Biobank: this research has been conducted using data from UK Biobank (www.ukbiobank.ac.uk). UK Biobank is generously supported by its founding funders the Wellcome Trust and UK Medical Research Council, as well as the Department of Health, Scottish Government, the Northwest Regional Development Agency, British Heart Foundation, and Cancer Research UK.

## Author contributions

L. Sun, W. Qin, C. Yu, and Y. He conceptualized the study. Y. He and C. Yu supervised the project. L. Sun, W. Qin, X. Liang, and Y. He designed the methodology. L. Sun, W. Qin, and X. Liang analyze the data. L. Sun and X. Liang developed visualizations. C. Yu, K. Li, F. Feng, C. Lin, Y. Yu, J. Qiu, C. Wang, L. Zhang, Xiaochu Zhang, H. Song, W. Men, Y. Duan, Z. Ye, P. Zhang, X. Fan, Q. Cai, S. Qiu, Q. Dong, S. Tao, M. Wang, Q. Gong, Y. Tian, P. Liang, Zeyu Liu, W. Zhu, Jintao Zhang, F. Xie, J. Feng, Jing Zhang, Chao Liu, Q. Qian, Bing Zhang, M. Meng, L. Hu, J. Gao, T. Jiang, X. Zhu, Y. He, Yuhan Zhang, Liping Liu, Hanjun Liu, W. Liao, D. Wang, H. Wang, T. Guo, Z. Dai, S. Lui, K. Xu, L. Li, P. Xie, C. Feng, G. Cui, J. Wu, X. Yin, G. Ding, J. Xian, L. Zhao, J. Lu, Zhifen Liu, Y. Han, Z. Yuan, Xilin Zhang, T. Si, F. Zhou, Y. Bi, D. Wu, F. Gao, Fei Wang, S. Qin, G. Wang, Feng Chen, Z. Zhang, J. Sui, Huafu Chen, J. Cai, S. Liu, Z. Geng, C. Zhang, N. Mao, H. Yin, B. Liu, H. Ma, B. Gao, Y. Miao, X. Kong, Y. Zhou, Li. Liu, J. Hu, L. Wang, Quan Zhang, H. Shu, Peijun Wang, T.M.C. Lee, Q. Cao, L. Yang, Xi Zhang, W. Luo, M. Liang, H. Yao, M. Li, H. Huang, Y. Peng, Z. Han, C. Zhou, H. Xu, M. Feng, W. Shen, Y. Hu, Huajun Chen, Ying Wang, G. Gong, Z. Yan, Xiaojun Xu, J. Liu, G. Chen, Pan Wang, Y. Yang, D. Yao, T. Han, Huiguang He, C. Chen, Q. Zou, Hesheng Liu, H. Zhang, C. Chai, C. Lu, Y. Tu, Yong Liu, D. Lin, W. Zhao, Xiufeng Xu, X. Liu, Z. Cui, Zheng Wang, R. Huang, Z. Li, Yunzhe Liu, X. Li, X. Yang, N. Zhang, A. Chen, Bin Zhang, P. Qin, Chen Liu, Z. Yao, Yanjun Wei, H. Yuan, Feng Wang, Yu Zhang, Quan Zhang, F. Hu, H. Xie, X. Wu, Jiaojian Wang, G. Fan, Zhiqun Wang, D. Zhang, H. Zhong, Yonggang Wang, L. Bai, Yongmei Li, X. Wei, Jinhui Wang, Yi Zhang, Hongjian He, S. Li, T. Zhang, F. Jiang, Jian Yang, Feiyan Chen, F. Liu, Huaigui Liu, N. Chen, Jinzhu Yang, B. Hou, C. Huang, J. Zhu, H. Cai, D. Wei, Q. Chen, Ying Wei, P. Miao, Yunxia Li, Yaou Liu, N. Yang, X. Gao, Yujie Liu, Y. Shen, X. Huang, and G. Ji collected a subset of the data for this study. L. Sun, X. Liang, and Y. He wrote the manuscript. All authors reviewed the final manuscript.

## Reference

1. Bethlehem, R.A.I., et al. Brain charts for the human lifespan. Nature 604, 525–533 (2022).

2. Rutherford, S. Charting brain growth and aging at high spatial precision. eLife 11, e72904 (2022).

3. Ge, R., et al. Normative modelling of brain morphometry across the lifespan with CentileBrain: algorithm benchmarking and model optimisation. Lancet Digit Health 6, e211–e221 (2024).

4. Sun, L., et al. Human lifespan changes in the brain’s functional connectome. Nat Neurosci 28, 891– 901 (2025).

5. Rutherford, S., et al. Evidence for embracing normative modeling. eLife 12, e85082 (2023).

6. Marquand, A.F., et al. Conceptualizing mental disorders as deviations from normative functioning. Mol Psychiatry 24, 1415–1424 (2019).

7. Segal, A., et al. Embracing variability in the search for biological mechanisms of psychiatric illness. Trends Cogn Sci 29, 85–99 (2025).

8. Tang, Y., et al. Brain structure differences between Chinese and Caucasian cohorts: A comprehensive morphometry study. Hum Brain Mapp 39, 2147–2155 (2018).

9. Liang, P., et al. Construction of brain atlases based on a multi-center MRI dataset of 2020 Chinese adults. Sci Rep 5, 18216 (2015).

10. Ge, J., et al. Increasing diversity in connectomics with the Chinese Human Connectome Project. Nat Neurosci 26, 163–172 (2023).

11. Dong, H.-M., et al. Charting brain growth in tandem with brain templates at school age. Sci Bul 65, 1924–1934 (2020).

12. Zhao, T., et al. Unbiased age-specific structural brain atlases for Chinese pediatric population. Neuroimage 189, 55–70 (2019).

13. Xie, W., Richards, J.E., Lei, D., Lee, K. & Gong, Q. Comparison of the brain development trajectory between Chinese and U.S. children and adolescents. Front Syst Neurosci 8, 249 (2014).

14. Fan, X.R., et al. A longitudinal resource for population neuroscience of school-age children and adolescents in China. Sci Data 10, 545 (2023).

15. Fu, J., et al. Cross-ancestry genome-wide association studies of brain imaging phenotypes. Nat Genet 56, 1110–1120 (2024).

16. Tooley, U.A., Bassett, D.S. & Mackey, A.P. Environmental influences on the pace of brain development. Nat Rev Neurosci 22, 372–384 (2021).

17. Northoff, S.H.a.G. Culture-sensitive neural substrates of human cognition: a transcultural neuroimaging approach. Nat Rev NeuroSci 9, 646–654 (2008).

18. Brouwer, R.M., et al. Genetic variants associated with longitudinal changes in brain structure across the lifespan. Nat Neurosci 25, 421–432 (2022).

19. Xu, J., et al. Global urbanicity is associated with brain and behaviour in young people. Nat Hum Behav 6, 279–293 (2022).

20. Rutherford, S. To which reference class do you belong? Measuring racial fairness of reference classes with normative modeling. arXiv (2024).

21. Li, J. Cross-ethnicity/race generalization failure of behavioral prediction from resting-state functional connectivity. Science Advance 8, eabj1812 (2022).

22. Desikan, R.S., et al. An automated labeling system for subdividing the human cerebral cortex on MRI scans into gyral based regions of interest. Neuroimage 31, 968–980 (2006).

23. Pini, L., et al. Brain atrophy in Alzheimer’s Disease and aging. Ageing Res Rev 30, 25–48 (2016).

24. Thompson, P.M. Dynamics ofGray Matter Loss in Alzheimer’s Disease. J Neurosci 23, 994–1005 (2003).

25. Young, P.N.E., et al. Imaging biomarkers in neurodegeneration: current and future practices. Alzheimers Res Ther 12, 49 (2020).

26. Segal, A., et al. Regional, circuit and network heterogeneity of brain abnormalities in psychiatric disorders. Nat Neurosci 26, 1613–1629 (2023).

27. Wolfers, T., et al. Mapping the Heterogeneous Phenotype of Schizophrenia and Bipolar Disorder Using Normative Models. JAMA Psychiatry 75, 1146–1155 (2018).

28. Dima, D., et al. Subcortical volumes across the lifespan: Data from 18,605 healthy individuals aged 3-90 years. Hum Brain Mapp 43, 452–469 (2022).

29. Frangou, S., et al. Cortical thickness across the lifespan: Data from 17,075 healthy individuals aged 3-90 years. Hum Brain Mapp 43, 431–451 (2022).

30. Porter, J.N., et al. Age-related changes in the intrinsic functional connectivity of the human ventral vs. dorsal striatum from childhood to middle age. Dev Cogn Neurosci 11, 83–95 (2015).

31. Tang, Y., et al. The construction of a Chinese MRI brain atlas: a morphometric comparison study between Chinese and Caucasian cohorts. Neuroimage 51, 33–41 (2010).

32. Peterson, R.E., et al. Genome-wide Association Studies in Ancestrally Diverse Populations: Opportunities, Methods, Pitfalls, and Recommendations. Cell 179, 589–603 (2019).

33. Gao, Y., et al. A pangenome reference of 36 Chinese populations. Nature 619, 112–121 (2023).

34. Xu, J., et al. Effects of urban living environments on mental health in adults. Nat Med 29, 1456–1467 (2023).

35. Gao, W., et al. A review on neuroimaging studies of genetic and environmental influences on early brain development. Neuroimage 185, 802–812 (2019).

36. Han, S. & Ma, Y. Cultural differences in human brain activity: a quantitative meta-analysis. Neuroimage 99, 293–300 (2014).

37. Obermeyer, Z. Dissecting racial bias in an algorithm used to manage the health of populations. Science 366, 447–453 (2019).

38. Martin, A.R. Clinical use of current polygenic risk scores may exacerbate health disparities. Nat Genet 51, 584–591 (2019).

39. Kamiza, A.B., et al. Transferability of genetic risk scores in African populations. Nat Med 28, 1163–1166 (2022).

40. Genomics is failing on diversity. Nature (2016).

41. Xu, Q., et al. CHIMGEN: a Chinese imaging genetics cohort to enhance cross-ethnic and cross-geographic brain research. Mol Psychiatry 25, 517–529 (2020).

42. Di Biase, M.A., et al. Mapping human brain charts cross-sectionally and longitudinally. Proc Natl Acad Sci U S A 120, e2216798120 (2023).

43. Glasser, M.F., et al. The minimal preprocessing pipelines for the Human Connectome Project. Neuroimage 80, 105–124 (2013).

44. Ji, J.L., et al. QuNex-An integrative platform for reproducible neuroimaging analytics. Front Neuroinform 17, 1104508 (2023).

45. Alfaro-Almagro, F., et al. Image processing and Quality Control for the first 10,000 brain imaging datasets from UK Biobank. Neuroimage 166, 400–424 (2018).

46. Hagler, D.J., Jr., et al. Image processing and analysis methods for the Adolescent Brain Cognitive Development Study. Neuroimage 202, 116091 (2019).

47. Wang, L., et al. iBEAT V2.0: a multisite-applicable, deep learning-based pipeline for infant cerebral cortical surface reconstruction. Nat Protoc 18, 1488–1509 (2023).

48. Makropoulos, A., et al. The developing human connectome project: A minimal processing pipeline for neonatal cortical surface reconstruction. Neuroimage 173, 88–112 (2018).

49. Williams, L.Z.J., et al. Structural and functional asymmetry of the neonatal cerebral cortex. Nat Hum Behav 7, 942–955 (2023).

50. Wu, Z., et al. Construction of 4D infant cortical surface atlases with sharp folding patterns via spherical patch-based group-wise sparse representation. Hum Brain Mapp 40, 3860–3880 (2019).

51. D. Mikis Stasinopoulos, R.A.R. Generalized Additive Models for Location Scale and Shape (GAMLSS) in R. J Stat Soft 23 (2007).

52. Borghi, E., et al. Construction of the World Health Organization child growth standards: selection of methods for attained growth curves. Stat Med 25, 247–265 (2006).

53. Rosen, A.F.G., et al. Quantitative assessment of structural image quality. Neuroimage 169, 407–418 (2018).

54. Dunn, P.K. & Smyth, G.K. Randomized Quantile Residuals. J Comput Graphical Stat 5, 236–244 (1996).

